# gMCSpy: Efficient and accurate computation of Genetic Minimal Cut Sets in Python

**DOI:** 10.1101/2024.02.02.578370

**Authors:** Carlos Javier Rodriguez, Naroa Barrena, Danel Olaverri-Mendizabal, Idoia Ochoa, Luis V. Valcarcel, Francisco J. Planes

## Abstract

**Motivation:** The identification of minimal genetic interventions that modulate metabolic processes constitutes one of the most relevant applications of genome-scale metabolic models (GEMs). The concept of Minimal Cut Sets (MCSs) and its extension at the gene level, genetic Minimal Cut Sets (gMCSs), have attracted increasing interest in the field of Systems Biology to address this task. Different computational tools have been developed to calculate MCSs and gMCSs using both commercial and open-source software.

**Results:** Here, we present *gMCSpy*, an efficient Python package to calculate gMCSs in GEMs using both commercial and non-commercial optimization solvers. We show that *gMCSpy* substantially overperforms our previous computational tool GMCS, which exclusively relied on commercial software. Moreover, we compared *gMCSpy* with recently published competing algorithms in the literature, finding significant improvements in both accuracy and computation time. All these advances make *gMCSpy* an attractive tool for researchers in the field of Systems Biology for different applications in health and biotechnology.

**Availability and Implementation:** The Python package *g*MCSpy can be accessed at: https://github.com/PlanesLab/gMCSpy

**Supplementary Information:** 

## Introduction

The large number of interrelated reactions that support life makes the characterization of biological systems a daunting task. Genome-scale metabolic models (GEMs) have emerged in the last two decades to address this complexity. In particular, GEMs provide a comprehensive representation of the metabolic and genetic interplay in an organism, aiming to offer a holistic view of cellular metabolism by integrating genomic, biochemical, and physiological information (Gu *et al*., 2019). Importantly, different molecular layers in GEMs are connected via gene-protein-reaction (GPRs) rules, which describe how genes translate into the enzymes of specific reactions that produce/consume metabolites. In recent years, the field of systems biology has witnessed significant advancements in the analysis of GEMs, with a particular focus on identifying potential intervention strategies for different clinical and biotechnological applications.

An influential concept for the identification of optimal intervention strategies in GEMs is Minimal Cut Sets (MCSs). MCSs define a minimal (irreducible) set of reactions whose deletion leads to a desired metabolic phenotype, e.g. infeasible biomass production or optimal production of a compound of biotechnological interest (Klamt and Gilles, 2004). We introduced a closely related concept called genetic Minimal Cut Sets (gMCSs), which define minimal intervention strategies at the gene level (Apaolaza *et al*., 2017). Different algorithms have been developed to calculate both MCSs and gMCSs in large GEMs (von Kamp and Klamt, 2014; Pratapa *et al*., 2015; Schneider *et al*., 2020). In particular, we developed a function in the COBRA toolbox (Heirendt *et al*., 2019) to carry out this task (Apaolaza *et al*., 2019), called here *GMCS*. More recently, *StrainDesign* was released (Schneider *et al*., 2022), a Python library that improves previous developments of the same group and extends their framework to an open-source platform.

Here, we present *gMCSpy*, a novel Python package that calculates gMCSs for GEMs. *gMCSpy* integrates several algorithmic improvements with respect to our previous tool, *GMCS*, which was built in MATLAB environment. Furthermore, *gMCSpy* allows the user to search for gMCSs with both commercial and open-source mixed-integer linear programming (optimization) solvers. We show that *gMCSpy* overperforms *GMCS* and *StrainDesign* in computation time and completeness of the solutions in a benchmark of 6 relevant GEMs. Overall, in our attempt to release an open-source framework, *gMCSpy* demonstrates computational and accuracy advances with respect to previous methods in the literature.

## Methods

*gMCSpy* is an open-source package written in Python to calculate gMCSs in GEMs. The package was built using *COBRApy* (Ebrahim *et al*., 2013) to conform with the standardization of GEMs and take advantage of model manipulations previously developed by the COBRA community.

In order to compute gMCSs, we implemented the Mixed Integer Linear Programming (MILP) model defined in the work of Apaolaza *et al*., 2019 (see *Supplementary Methods*). *gMCSpy* includes a common interface to define this MILP model and translation functions to compute gMCSs with different solvers, namely the commercial solvers Gurobi (Gurobi Optimization LLC, 2023) and IBM ILog CPLEX (Cplex, 2009), and the open-source solver SCIP (Bestuzheva *et al*., 2023). This constitutes an advance with respect to our previous tool, GMCS, which was developed in MATLAB environment exclusively for CPLEX. Moreover, *gMCSpy* can use the latest versions of CPLEX, currently V22.1.0, in contrast, the latest version of CPLEX compatible with MATLAB is V12.10.0.

A critical part in our methodology is the computation of matrix G, which defines the gene knockout constraints in our MILP model (see *Supplementary Methods*). This step is performed by the function *buildGMatrix*, which has been substantially improved in *gMCSpy* with respect to our previous work (Apaolaza *et al*., 2019). In particular, *buildGMatrix* requires the function *parseGPRToModel*, which transforms GPR rules into artificial reaction networks (GPR networks) (Apaolaza *et al*., 2019; Barrena *et al*., 2023). We have implemented an efficient recursive strategy in Python to reduce the computational expenditure in constructing GPR networks. This function allows us to efficiently deal with complex GPR rules, in contrast to other methods that limit the size of GPRs that they can handle (Schneider *et al*., 2022). Importantly, we have also refined the definition of matrix G to more accurately enumerate gMCSs in increasing length order (see *Supplementary Methods*).

The main function to calculate gMCSs in *gMCSpy* is *calculateGeneMCS*. It can be used to perform a global search (with all genes) but also to identify gMCSs involving a specific subset of genes, as done in Apaolaza *et al*., 2017, with the targetKOs attribute. Likewise, we can compute nutrient-genetic MCSs (ngMCSs), interventions that combine nutrient deprivations in the environment and gene knockouts (Apaolaza *et al*., 2022), using the *isNutrient* attribute. The same analysis can be done at the reaction level with the function *calculateMCS*. Different examples are available in the *gMCSpy* documentation to illustrate these functions.

## Results

We compared *gMCSpy* with two published tools in the literature: our previous MATLAB tool, GMCS, and StrainDesign. *gMCSpy* and *StrainDesign* were benchmarked here with two commercial solvers: CPLEX and Gurobi; while *GMCS* was examined only with CPLEX, as it is not designed to work with Gurobi. The results obtained with the open-source solver, SCIP, can be found in *Supplementary Figure 1*. For this side-by-side comparison, we used *E. coli* core (Orth *et al*., 2010) and the most recent GEMs of *E. coli*, iML1515 (Monk *et al*., 2017); *P. putida*, iJN1463 (Nogales *et al*., 2020); *S. cerevisiae*, Yeast-GEM v8.7.0 (Lu *et al*., 2019); and human cells, Human-GEM v1.16.0 (Robinson *et al*., 2020) (Table 1). In the case of human cells, we considered two cases: under the most general growth medium (Human-GEM v1.16.0) and under Ham*’*s growth medium (Human-GEM v1.16.0_CultureMedia). Using the RAVEN toolbox, we applied the Ham*’*s growth medium and removed all reactions and associated metabolites that cannot undertake any flux in this setting (Wang *et al*., 2018). We assessed the capacity of the different approaches to extract all single, double and triple gene deletions which result to be lethal for proliferation, as well as the required computation time. The benchmark was performed in an Intel Xeon Gold 8248R using 16 threads and 64GB of RAM. Full details can be found in Figure 1.

**Table 1:** Summary of GEMs used in the benchmarking study of gMCS computation. We considered E. coli core (Orth *et al*., 2010); E. coli, iML1515 (Monk *et al*., 2017); P. putida, iJN1463 (Nogales *et al*., 2020); S. cerevisiae, Yeast-GEM v8.7.0 (Lu *et al*., 2019); and *human* cells, Human-GEM v1.16.0 (Robinson *et al*., 2020), respectively. In the case of human cells, we considered two cases: under the most general growth medium (Human-GEM v1.16.0) and under Ham*’*s growth medium (Human-GEM v1.16.0_CultureMedia). The dimension of each case considered, in terms of number of reactions, metabolites and genes, is provided. The last three columns analyze whether (or not) the different methods considered, the considered methods can be applied to search for gMCSs for the corresponding GEM.

| Model | No. Reactions | No. Metabolites | No. Genes | Gmcspsy | StrainDesign | GMCS |
| --- | --- | --- | --- | --- | --- | --- |
| *E. coli core* | 95 | 72 | 137 | ✓ | ✓ | ✓ |
| *iML1515* | 2712 | 1877 | 1516 | ✓ | ✓ | ✓ |
| *iJN1463* | 2927 | 2153 | 1462 | ✓ | ✓ | ✓ |
| *Yeast-GEM* | 4131 | 2806 | 1163 | ✓ | × | ✓ |
| *Human-GEM v1.16.0* | 11944 | 7118 | 2897 | ✓ | × | ✓ |
| *Human-GEM v1.16.0_CultureMedia* | 11509 | 6800 | 2897 | ✓ | × | ✓ |

**Figure 1:**
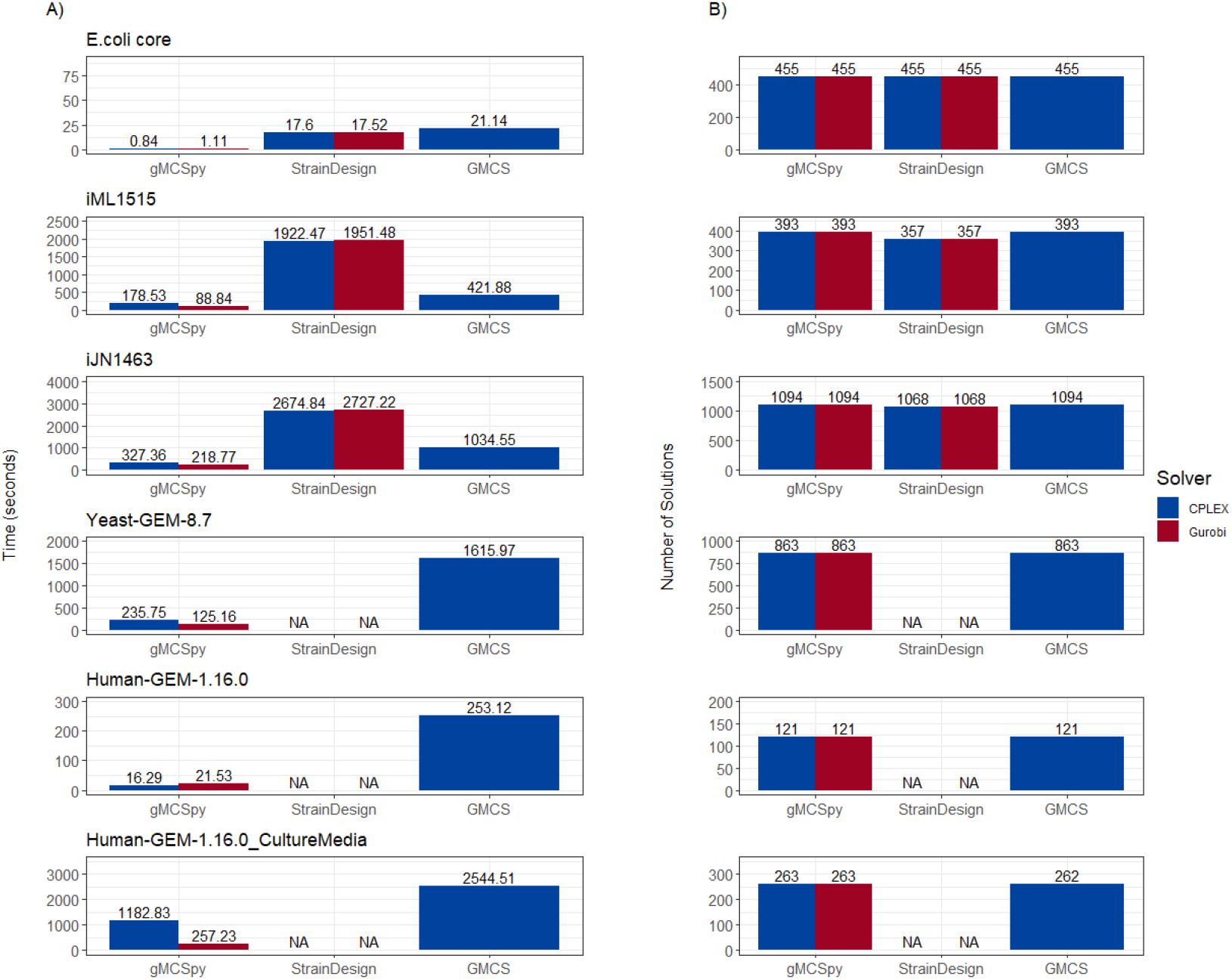
Benchmark of existing tools to compute gMCSs in GEMs. A) Computation times to calculate gMCS up to length 3. *i.e*., lethal single, double, and triple knockouts, for each of the cases analyzed with *gMCSpy*, StrainDesign and GMCS. Mean values across 10 different runs are shown at the top of bars. *‘*NA*’* means that the calculation was not possible. B) Number of gMCSs up to length 3 that were found for each of the cases analyzed with *gMCSpy*, StrainDesign and GMCS.

First, we note that not all GEMs were compatible with StrainDesign (Table 1). This incompatibility comes from GPRs, as *StrainDesign* cannot handle GPRs larger than a certain length. This implies that large GPRs need to be removed before using *StrainDesign*, in contrast to *gMCSpy* or GMCS, which can be directly used to search for gMCSs.

Second, *gMCSpy* substantially reduces the computation time with respect to *StrainDesign* in all the GEMs tested (Figure 1A). In some of the scenarios, we obtained improvements of nearly 20-fold increase in speed with respect to S*trainDesign* (e.g., with Gurobi in iML1515). Moreover, *gMCSpy* obtained a better performance in Gurobi than in CPLEX in most cases. This is not the case in *StrainDesign*, where both solvers had a similar behavior. Importantly, *StrainDesign* failed to recover all the solutions in some GEMS. For example, it could not recover 36 lethal triple gene knockouts in iML1515. This was not the case in *gMCSpy*, which recovered all the solutions.

With respect to our previous tool, *GMCS*, we note several performance improvements. First, as mentioned in the Methods section, a key improvement of *gMCSpy* over *GMCS* is related to the computation of matrix G. We observed significant differences in computation time (two-sided Wilcoxon test p-value = 0.03125), finding drastic improvements in some of the scenarios (*Supplementary Figure 2*). For example, *gMCSpy* took 3.94 seconds to build matrix G in Human-GEM v1.16.0_CultureMedia while GMCS took 227.97 seconds. Second, we slightly modified matrix G to avoid the case found in Human-GEM v1.16.0_CultureMedia, where one lethal triple gene knockout was missed with GMCS (Figure 1B) (see *Supplementary Methods*). This modification in our approach has a negligible impact on the computation performance of *gMCSpy* (Supplementary Figure 3). Overall, we show superior performance across all models in both solvers, presenting more than 10-fold reduction in computation time in some scenarios tested (*e*.*g*. Yeast-GEM-8.7 in Gurobi). These results clearly show that *gMCSpy* is more efficient and accurate in computing gMCSs than our previous MATLAB tool, *GMCS*.

## Discussion

The capacity of predicting key genetic interventions has profound implications for various areas in health and biotechnology. The gMCS framework offers a valuable tool for optimizing interventions within biological systems, particularly in the field of drug discovery, where it can aid in the identification of potential drug targets that modulate cellular metabolism in different disease areas, opening new avenues for the development of personalized medicine.

*gMCSpy* constitutes an efficient open-source Python package to calculate gMCSs in GEMs using both commercial and non-commercial optimization solvers. It substantially improves our previous tool, *GMCS*, which was developed in MATLAB environment and limited to CPLEX, as shown in the benchmarking study summarized in Figure 1. Moreover, *gMCSpy* overperforms in both accuracy and computation time *StrainDesign*, a competing algorithm in the literature. All these advances make *gMCSpy* an attractive tool for researchers in the field of Systems Biology.

## Acknowledgements

This work was supported by the Minister of Economy and Competitiveness of Spain [PID2019-110344RB-I00 and PID2022-143298OB-I00, F.J.P., and PID2021-126718OA-I00, I.O.], PIBA Programme of the Basque Government [PIBA_2020_01_0055, F.J.P.], ERANET program ERAPerMed [MEET-AML, F.P.], Ramon y Cajal contract [RYC2019-028578-I, I.O.], Gipuzkoa Fellows grant [2022-FELL-000003-01, I.O], Elkartek programme of the Basque Government [KK-2022/00045, F.J.P., KK-2023/00001, I.O.], Ramon Areces grant [to FJP]. N.B. received his salary from a Basque Government predoctoral grant [PRE_2021_2_0025]. The funders had no role in study design, data collection and analysis, decision to publish, or preparation of the manuscript.

