## Supplemental Material for "gMCSpy: Efficient and accurate computation of Genetic Minimal Cut Sets in Python"

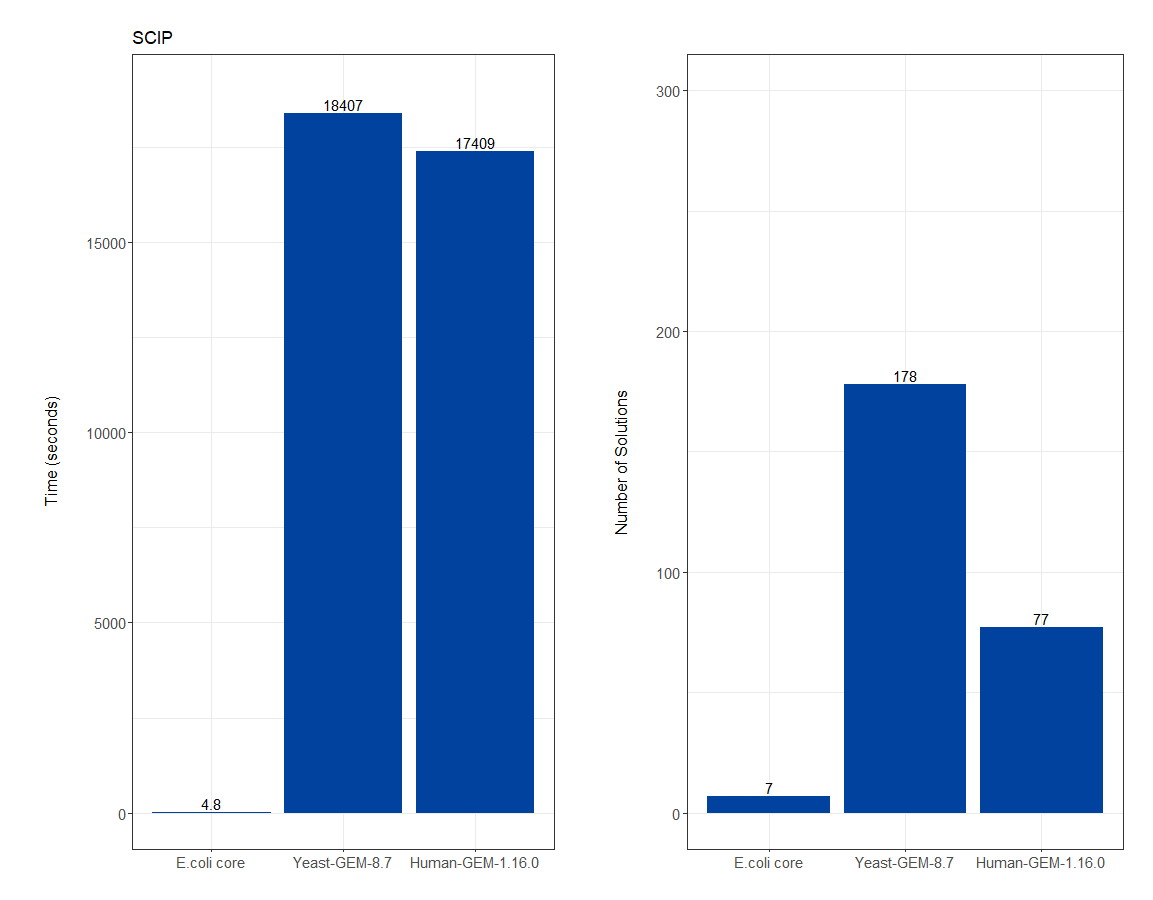

**Supplementary Figure 1: SCIP performance with gMCSpy.** We present the performance of gMCSpy using the optimization solver SCIP (Bestuzheva et al., 2023) for 3 different GEMs: E. coli core (Orth et al., 2010), Yeast-GEM v8.7.0 (Lu et al., 2019) and Human-GEM v1.16.0 (Robinson et al., 2020). Results are shown for the computation of lethal single gene knockouts. We fixed a time limit of 10,000 seconds per solution. gMCSpy found all the solutions in E. coli core and Yeast-GEM v8.7.0, but it failed to recover the whole set of solutions in Human-GEM and found only 77 out 92 interventions. The difference in computation time with respect to Gurobi and CPLEX is very relevant.

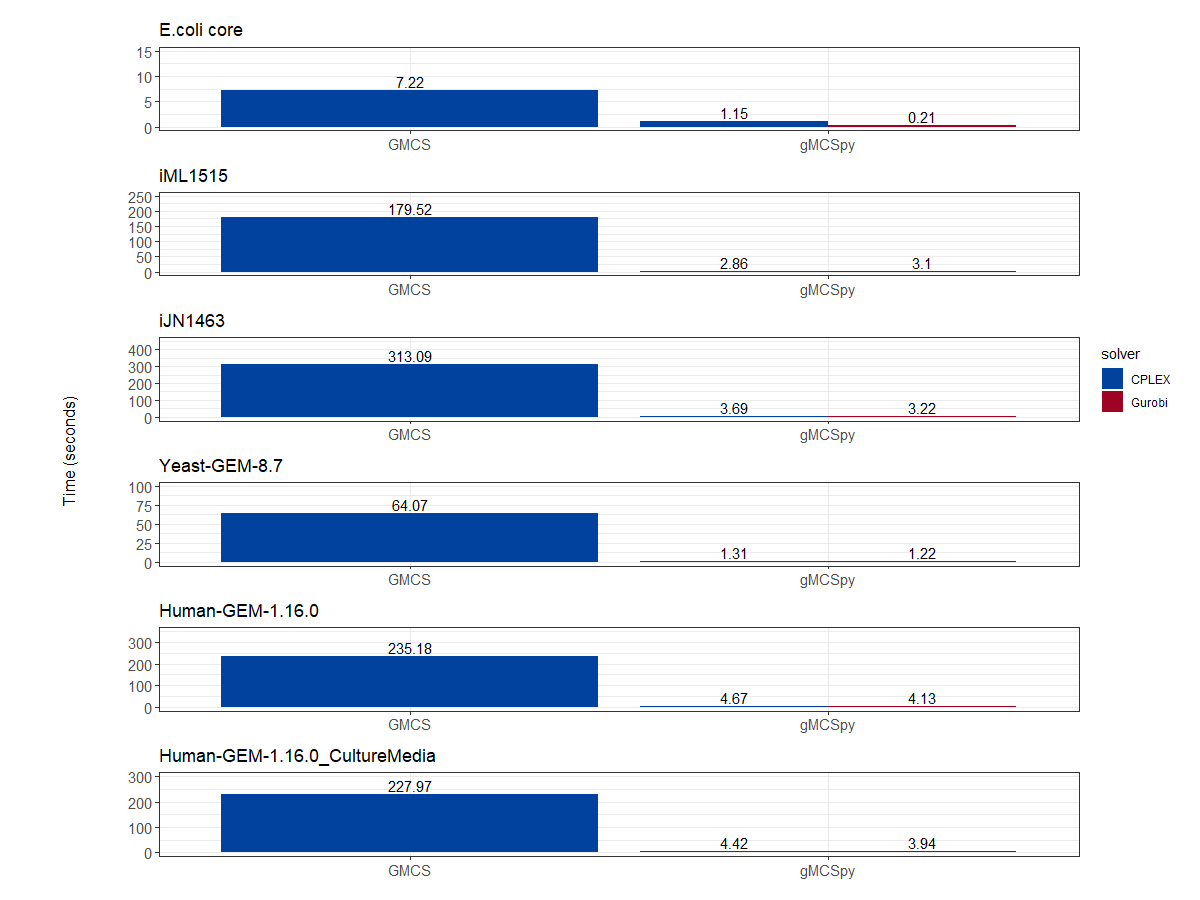

**Supplementary Figure 2: Computation time for the calculation of matrix G with gMCSpy and GMCS.** We considered in this study E. coli core (Orth et al., 2010) and the most recent GEMs of E. coli, P. putida, S. cerevisiae and human cells: iML1515 (Monk et al., 2017), iJN1463 (Nogales et al., 2020), Yeast-GEM v8.7.0 (Lu et al., 2019) and Human-GEM v1.16.0 (Robinson et al., 2020), respectively. In the case of human cells, we considered two cases: under the most general growth medium (Human-GEM v1.16.0) and under Ham’s growth medium (Human-GEM v1.16.0_CultureMedia). For each GEM, we calculated the G matrix with gMCSpy and GMCS and measured the execution time (in seconds) with the 2 commercial solvers: CPLEX and Gurobi.

**Supplementary Figure 3: Performance of gMCSpy with different methodologies.** As noted in the main text, we implemented in gMCSpy the Mixed Integer Linear Programming (MILP) model defined in the work of Apaolaza et al., 2019, with a slight modification in the definition of matrix G that allows us to more accurately enumerate gMCSs in increasing length order. We compared here the performance of gMCSpy with our previous methodology (gMCSpy_old_MILP) and our new methodology (gMCSpy) with the two commercial solvers: CPLEX and Gurobi. Again, we considered E. coli core (Orth et al., 2010) and the most recent GEMs of E. coli, P. putida, S. cerevisiae and human cells: iML1515 (Monk et al., 2017), iJN1463 (Nogales et al., 2020), Yeast-GEM v8.7.0 (Lu et al., 2019) and Human-GEM v1.16.0 (Robinson et al., 2020), respectively. In the case of human cells, we considered two cases: under the most general growth medium (Human-GEM v1.16.0) and under Ham’s growth medium (Human-GEM v1.16.0_CultureMedia). A) Computation times (in seconds) to calculate gMCS up to length 3. i.e. lethal single, double and triple knockouts, for each of the cases analyzed with gMCSpy and gMCSpy_old_MILP. Mean values across 10 different runs are shown at the top of bars. B) Number of gMCSs up to length 3 that were found for each of the cases analyzed with gMCSpy and gMCSpy_old_MILP.

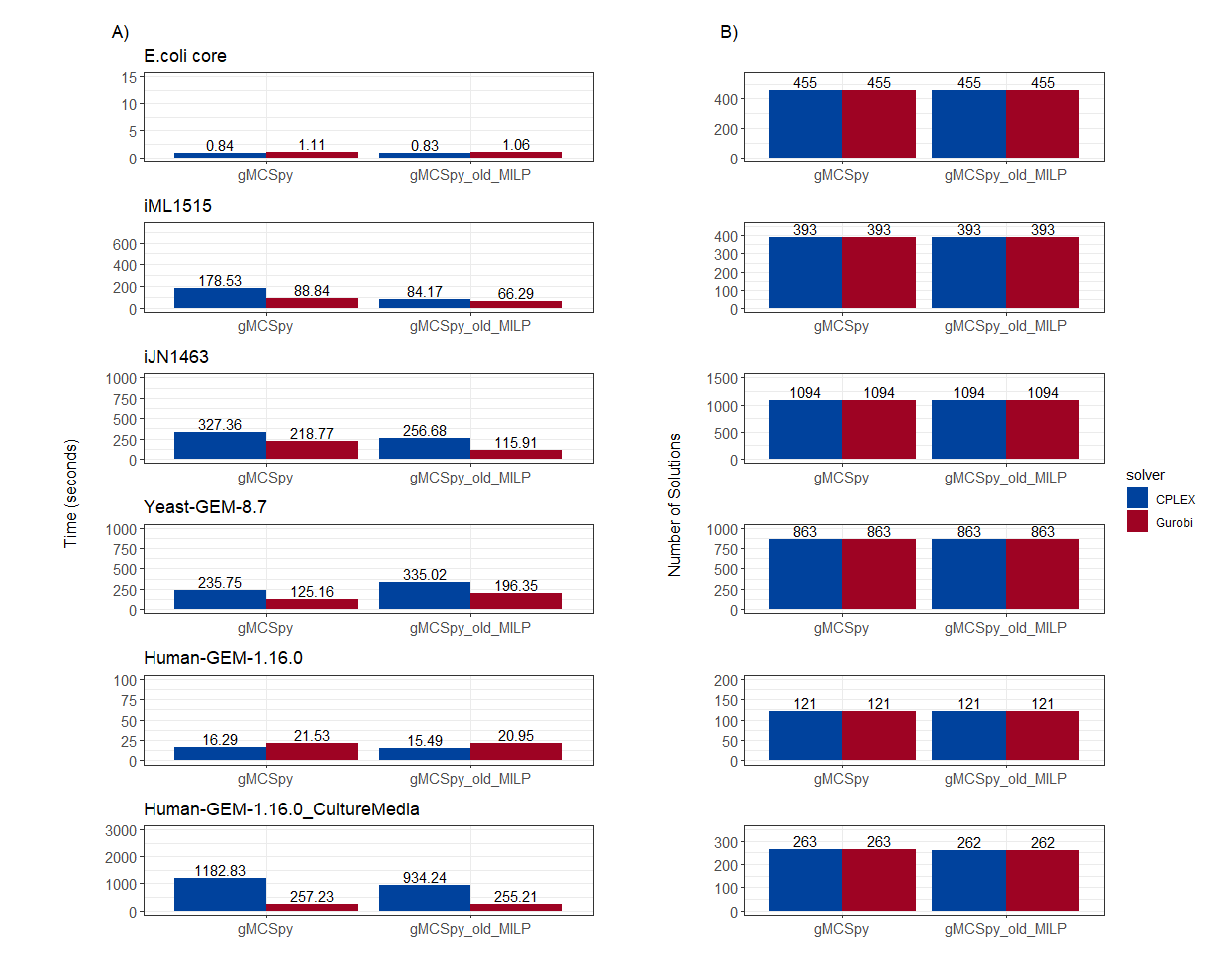

**Supplementary Methods**

**MILP formulation for the calculation of gMCSs**

In this section, we present a brief description of the mixed integer linear programming (MILP) model, previously developed in different works of our group (Apaolaza et al., 2017, 2019; Barrena et al., 2023), which allows us to search for genetic minimal cut sets (gMCS) in genome-scale metabolic models (GEMs).

For a particular GEM, the full set of reactions is typically represented with the stoichiometric matrix, here denoted as $S$, of dimensions $m\times n$, where $m$ and $n$ are the total number of metabolites and reactions, respectively. The values of each column represent the stoichiometric coefficients of the different metabolites in a particular reaction. Products and substrates in a reaction take positive and negative coefficients, respectively. Each reaction can carry a flux $r$; however, we do no not allow negative fluxes and, thus, each reversible reaction must be split into two irreversible reactions:

$r \geq0$ (Eq.1)

Considering the steady state and the mass balance equation, we can establish that consumption and production fluxes must be equal to zero:

$S\cdot r=0$ (Eq.2)

Next, we define the target metabolic task that we aim to block, typically the biomass reaction. Thus, we force a positive flux through this target task:

$t^{T}\cdot r \geq t^{*}$ (Eq.3)

where $t^{T}$represents a row vector in which all its elements are zero except for the reactions implied in the task; and $t^{*}$ is a positive constant.

Eq. 4 defines the gene knockout constraints. As detailed in the next section, each row in G specifies the subset of reactions blocked as a result of a minimal subset of gene knockouts, stored in *F* matrix.

$G\cdot r \leq0$ *(Eq.4)*

Eqs. 1-4 define the primal linear programming problem. Due to the inconsistency between Eq.3 and Eq.4, the primal problem is infeasible. In that situation, its associated dual problem defines an unbounded polyhedral cone, whose extreme rays represent minimal subsets of constraints in the primal problem that are inconsistent and lead to the underlying infeasibility. This is closely related with the concept of gMCSs, which are defined by Eq.3 and minimal subsets of gene knockout constraints in Eq.4. Accordingly, gMCSs can be calculated using the following MILP:

$minimize\sum_{i=1}^{i=l} d_{i}\cdot z_{i}$ (Eq.5)

$$s.t.$$

$N\cdot\left( \begin{matrix} u \\ v \\ w \end{matrix} \right)=\left[ S^{T} G^{T}-t \right]\cdot\left( \begin{matrix} u \\ v \\ w \end{matrix} \right)\geq0$ (Eq.6)

$\alpha\cdot z\leq v\leq M\cdot z$	(Eq.7)

$r^{*}\cdot w\leq-c, c>0$ (Eq.8)

$z_{\delta}\geq z_{\beta} \forall\left( \delta, \beta\right)| F\left( \beta\right)\supset F\left( \delta\right)$ (Eq.9)

$\sum_{i=1}^{i=l} z_{i}^{j}z_{i}\leq\sum_{i=1}^{i=l} z_{i}^{j}-1$ (Eq.10)

$v\geq0;w\geq0$ (Eq.11)

$u\in R^{m}, v\in R^{l}, w\in R, z \in B^{l}$ (Eq.12)

where $u$, $v$, and $w$ denote dual variables linked to the mass balance equation, gene knockout constraints, and the target (metabolic task) constraint, respectively. Eq. 6 defines the constraints associated to the dual problem. In addition, $z$ variables are binary variables associated with $v$ variables through Eq.7, which guarantees that if $z=1$ then $v>\alpha$ or if $z=$ 0 then $v=0$. Eq. 8 forces $w$ to be non-zero, incorporating the target constraint, Eq.3, into the infeasible primal problem. Eq.9 addresses the dependencies between dual variables $v$, which may yield non-minimal solutions. Finally, Eq.10 eliminates previous solutions ($z^{j}$) from the solution space and enables the enumeration of gMCSs in increasing number of gene knockout. Eqs. 11-12 define the nature of variables used in our MILP. The objective function, Eq.5, is discussed below.

**Calculation of matrix G**

In practice, before formulating the MILP problem, we need to define matrix G, which is required for the construction of the gene knockout constraints in Eq. 4. Each row in the G matrix is linked to a minimal subset of gene knockouts that disables a specific subgroup of reactions. In *gMCSpy*, for the definition of matrix G, we use a Python dictionary, data structure composed by key-value pairs, where each key must be unique and associated with a specific value. In our implementation, keys represent a subset of genes and values a list of reactions. We call the resulting Python dictionary *GDict*.

For illustration, consider the example network depicted in Supplementary Figure 4A, which involves 6 reactions, 4 metabolites and 7 genes in GPR rules. For the construction of *GDict*, we first focus on reactions controlled only by one gene, in our case $r_{1} \mathrm{and} r_{5}$. Next, we consider reactions that are only controlled by AND relationships followed by reactions with only OR relationships (only $r_{4}$ in the toy example). Finally, we deal with more complex GPRs, such as $r_{2} \mathrm{and} r_{3}$. For it, we first convert GPRs into Boolean trees using *COBRApy* (Ebrahim et al., 2013). The, we apply an efficient recursive function (developed in gMCSpy) that transforms Boolean trees into artificial reaction networks (GPR networks), as shown in Supplementary Figure 4B. We then search for MCSs that disrupts the target reaction in these GPR networks using the method described in Apaolaza et al., 2019, based on Minimal Cut Sets.

**Calculation of G matrix**

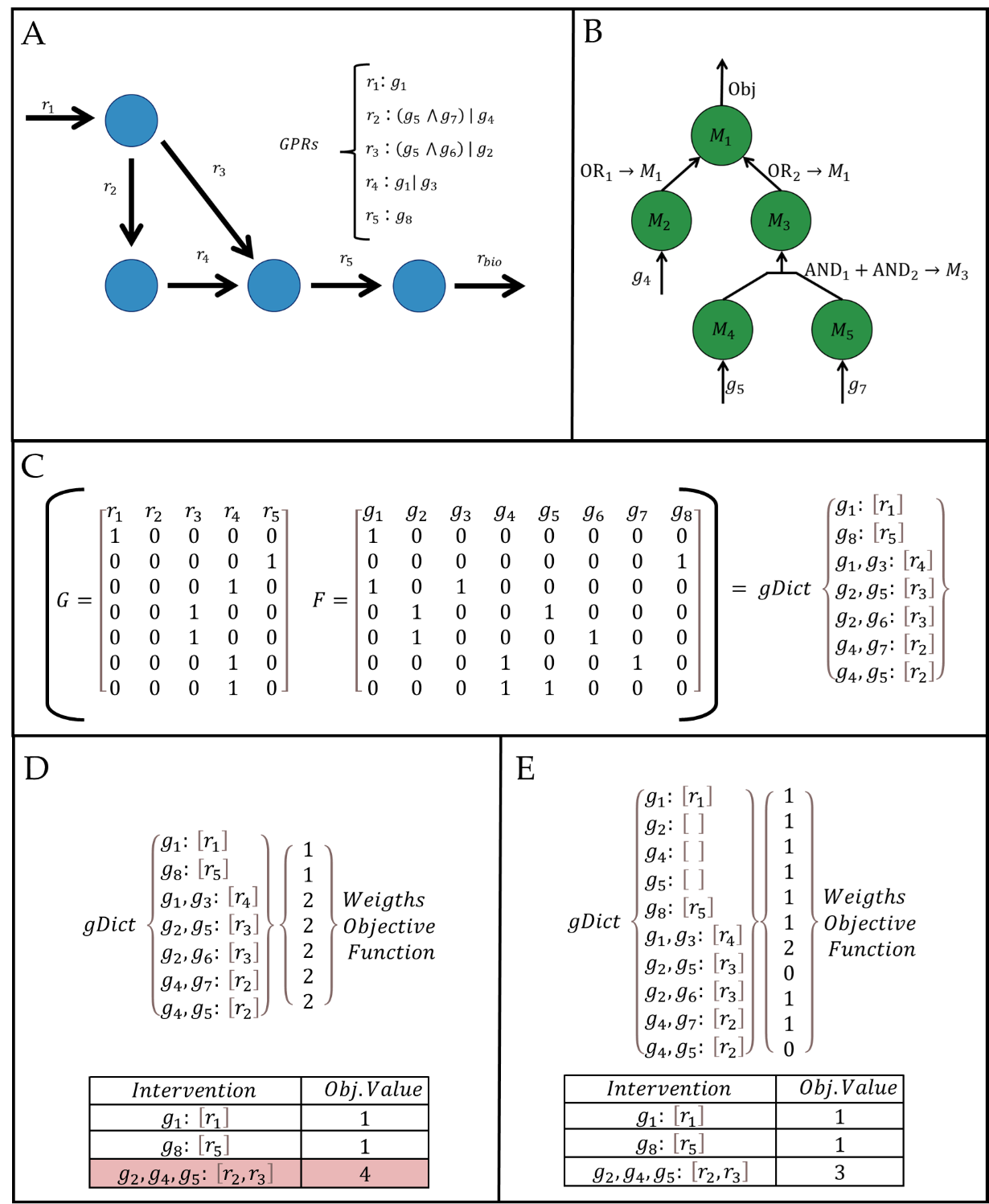

**Supplementary Figure 4: Toy example to illustrate the calculation of G matrix.** A) Example metabolic network involving 6 six reactions: r_1_, r_2_, r_3_, r_4_, r_5_ and r_bio_, and 8 genes: g_1_, g_2_, g_3_, g_4_, g_5_, g_6_, g_7_ and g_8_ in GPR rules; B) Example GPR network for r_2_. GPR networks are built using an efficient recursive function using Boolean tress available in COBRApy; C) Resulting G and F matrices, as well as data structure exemplifying the equivalent GDict for our toy example. D) One example of the expansion process between interventions, if the intersection of two keys yields a non-existing key in the GDict it is added as a null row. E) The equivalent GDict in methodology presented in Apaolaza et al. 2019 and associated weights in the objective function. In the toy example, the intervention $\{g_{2}, g_{4}, g_{5}\}$ would NOT be discovered when computing all single, double and triple knockouts. E) The GDict generated in gMCSpy inserting intersecting genes mitigates this eventuality increasing the accuracy of our methodology.

The resulting *GDict* in our toy example can be observed in Supplementary Figure 4C. *GDict* condenses all the information required to build Eq.4 and Eq.6 in our MILP into a single data structure, in contrast with our previous MATLAB approach requiring the calculation of matrices G and F, also shown in Supplementary Figure 4C. Note here that each row in G defines a specific subset of reactions blocks for a minimal subset of gene knockouts, which are defined in its associated row in F.

**Preprocessing expansion of interventions**

The objective function in our MILP, Eq.5, aims to minimize the number of gene knockouts to block the target metabolic task, according to a weighting system of $z$ variables that considers their dependencies, summarized in Eq.9. For illustration, in our toy example, the intervention associated with the first row in G and F, ${\{g}_{1}{:[r}_{1}]\}$, is contained in the intervention associated with the sixth row in G and F, ${\{g}_{1},{g_{3}:[r}_{4}]\}$. Thus, we need to implement the following constraint: $z_{1}-z_{6}\geq0$, which means that if we activate the intervention of the sixth row, $z_{6}=1$ (knockout of $g_{1}$and $g_{3}$), then the intervention of the first row must be also active, $z_{1}=1$ (knockout of $g_{1}$).

As detailed in Apaolazaet al., 2019, these dependency constraints generate an imbalance in the objective function that requires adjusting the weights of each binary variable $z_{i}$ in order to return exactly the number of genes involved in the output gMCS. In our previous methodology (GMCS), the weight of each variable $z_{i}$ is defined as the number of genes involved in its associated intervention minus the number of genes involved in other dependent interventions of smaller length. For example, the weight of dual variable $z_{6}$ in the objective function is $d_{6}=2-1=1$, since we need to subtract the number of genes in its dependent intervention, $z_{1}$. The resulting weights for our toy example are shown in Supplementary Figure 4D.

The caveat of this methodology is that we may find situations where the objective does not coincide exactly with the number of gene knockouts. This explains why one of the gMCSs of length 3 in the case of *Human-GEM v1.16.0_CultureMedia* is missed, as summarized in Figure 1B in the main text. Supplementary Figure 4D also shows one of the resulting gMCSs, {g_2_,g_4_,g_5_}, which incorrectly has an objective value of 4 and should be 3.

To address this issue, *gMCSpy* uses a different approach. In particular, we build a slightly modified G matrix and, therefore, *GDict*, as shown in Supplementary Figure 4E for our toy example. In particular, a null row is added to G for every gene that takes part in F matrix in more than one row but not as a single gene. This is the case in our toy example of the following genes: *g_2_, g_4_* and *g_5_*. For example, in the case of *g_5_*, it participates in two rows of F, {*g_2_, g_5_*} and {*g_4_, g_5_*}, but none of them as a single. This strategy requires a slightly larger G matrix and, thus, a higher number of dual variables. However, it corrects the weights of dual variables (Supplementary Figure 4E), guaranteeing that all solutions are recovered with the exact length of gene knockouts with a similar computational cost (Supplementary Figure 4E).
